# A pandemic clonal lineage of the wheat blast fungus

**DOI:** 10.1101/2022.06.06.494979

**Authors:** Sergio M. Latorre, Vincent M. Were, Andrew J. Foster, Thorsten Langner, Angus Malmgren, Adeline Harant, Soichiro Asuke, Sarai Reyes-Avila, Dipali Rani Gupta, Cassandra Jensen, Weibin Ma, Nur Uddin Mahmud, Md. Shåbab Mehebub, Rabson M. Mulenga, Abu Naim Md. Muzahid, Sanjoy Kumar Paul, S. M. Fajle Rabby, Abdullah Al Mahbub Raha, Lauren Ryder, Ram-Krishna Shrestha, Suwilanji Sichilima, Darren M. Soanes, Pawan Kumar Singh, Alison R. Bentley, Diane G. O. Saunders, Yukio Tosa, Daniel Croll, Kurt H Lamour, Tofazzal Islam, Batiseba Tembo, Joe Win, Nicholas J. Talbot, Hernán A. Burbano, Sophien Kamoun

**Affiliations:** Centre for Life’s Origins and Evolution, Department of Genetics, Evolution and Environment, University College London; London, UK; The Sainsbury Laboratory, University of East Anglia; Norwich Research Park, Norwich, UK; Graduate School of Agricultural Science, Kobe University; Kobe, Japan; Institute of Biotechnology and Genetic Engineering, Bangabandhu Sheikh Mujibur Rahman Agricultural University; Gazipur, Bangladesh; John Innes Centre, Norwich Research Park, Norwich, UK; Zambia Agricultural Research Institute, Mt. Makulu Central Research Station; Lusaka, Zambia; University of Exeter; Exeter, UK; International Maize and Wheat Improvement Center, (CIMMYT); Texcoco, Mexico; Laboratory of Evolutionary Genetics, Institute of Biology, University of Neuchâtel; Switzerland; Department of Entomology and Plant Pathology, University of Tennessee; Knoxville, USA

## Abstract

Wheat, the most important food crop, is threatened by a blast disease pandemic. Here, we show that a clonal lineage of the wheat blast fungus recently spread to Asia and Africa following two independent introductions from South America. Through a combination of genome analyses and laboratory experiments, we show that the decade-old blast pandemic lineage can be controlled by the Rmg8 disease resistance gene and is sensitive to strobilurin fungicides. However, we also highlight the potential of the pandemic clone to evolve fungicide-insensitive variants and sexually recombine with African lineages. This underscores the urgent need for genomic surveillance to track and mitigate the spread of wheat blast outside of South America, and to guide pre-emptive wheat breeding for blast resistance.

## MAIN TEXT

Plant disease outbreaks threaten the world’s food security at alarming levels (Fisher et al. 2020; Ristaino et al. 2021). Understanding pathogen evolution during epidemics is essential for developing a knowledge-based disease management response. Genomic surveillance meanwhile adds a unique dimension to coordinated responses to infectious disease outbreaks and is central to the Global Surveillance System (GSS) recently proposed to increase global preparedness to plant health emergencies (World Health Organization 2022-2032).

In wheat, yield losses caused by pests and diseases average over 20% (Savary et al. 2019). Wheat is currently threatened by the expanding blast pandemic caused by the ascomycete fungus *Magnaporthe oryzae* (Syn. *Pyricularia oryzae*), a formidable and persistent menace to major grain cereals that can cause total crop failure (Mehrabi and Ramankutty 2019). The disease first appeared in 1985 in Brazil but has been reported in Bangladesh and Zambia over the last years. The occurrence of wheat blast on three continents with climatic conditions highly conducive to its spread, is unprecedented and represents a very significant threat to global food security which is exacerbated by the unprecedented twin challenge of climate change and armed conflicts in major agricultural regions.

Wheat blast emerged in Brazil in 1985 following deployment of rwt3 wheat genotypes, which facilitated host jumps of *M. oryzae* from *Lolium* spp. to wheat through loss of function mutations in the PWT3 effector (Inoue et al. 2017). A lineage of the wheat blast fungus that first emerged in Bangladesh in 2016 was previously traced to the genetically diverse South American population (Islam et al. 2016). However, the genetic identity and origin of the causal agent of an African outbreak, first detected in Zambia in 2018, remains unknown (Tembo et al. 2020).

To determine the relationship between African wheat blast isolates from Zambia with populations from South America and Bangladesh, we selected 84 single nucleotide polymorphisms (SNPs) mined from the sequence data of Islam et al. (Islam et al. 2016) to discriminate between the Bangladesh lineage from other *M. oryzae* genotypes. We genotyped 537 *M. oryzae* samples from different geographical regions and hosts based on multiplex amplicon sequencing (MonsterPlex; see material and methods) (N=237) and publicly available genomes (N=351) (Fig. 1, Fig. S1, Table S1). The Zambian isolates (N=13, 2018-2020) are identical for the 84 SNPs to wheat isolates from Bangladesh (N=81, 2016-2020) and one genotype B71 from South America (Bolivia, N=1, 2012). We conclude that this “B71 lineage”, which emerged in Asia in 2016 and traces its origins to South America, is now established in Zambia.

**Fig. 1.**
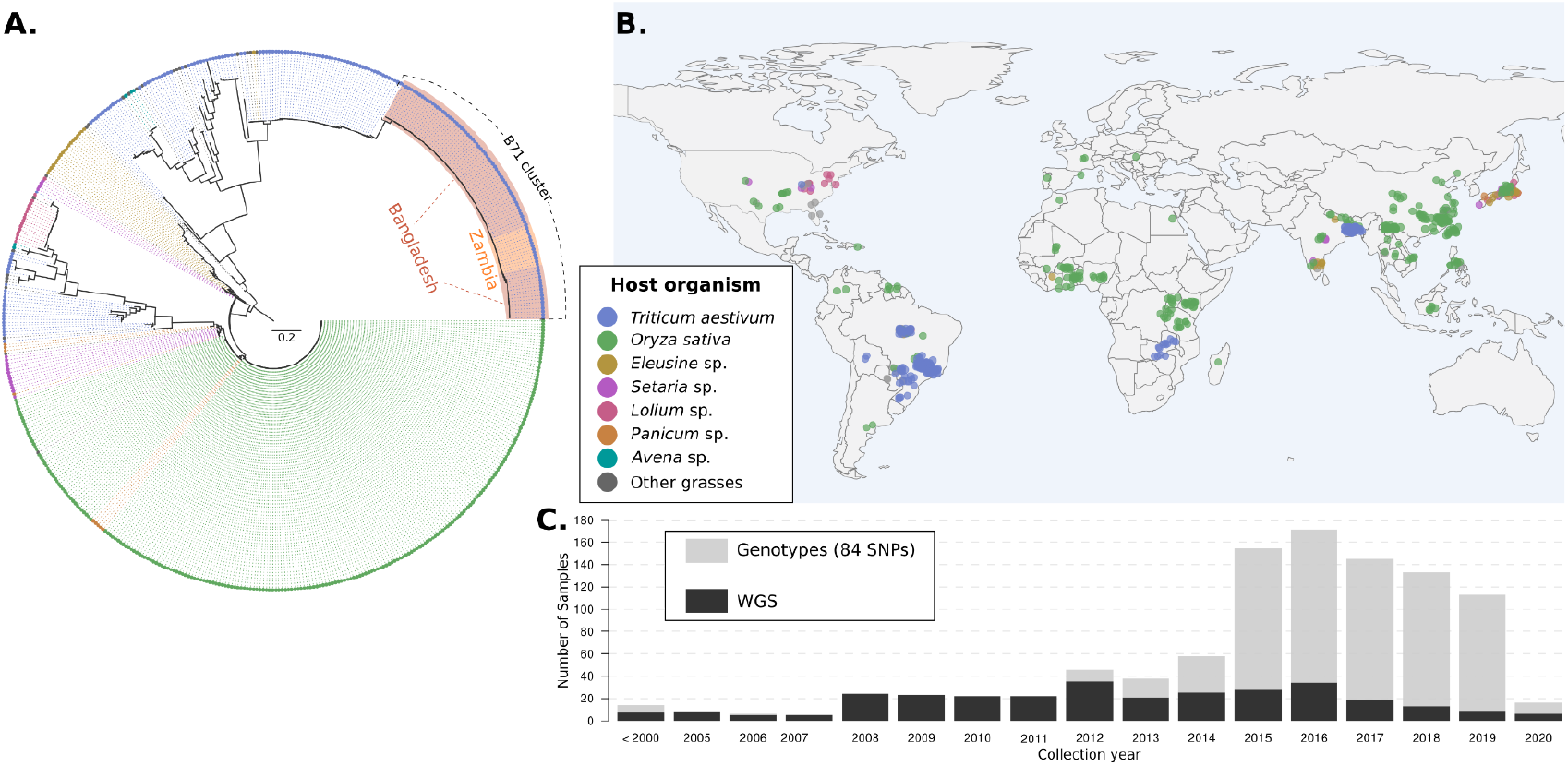
The emergence of wheat blast in Bangladesh and Zambia was caused by the B71 genetic lineage of *Magnaporthe oryzae*. **(A)** Neighbor-joining tree of 532 worldwide distributed *M. oryzae* isolates based on 84 concatenated SNPs obtained by multiplex amplicon sequence and/or genome sequences. The topology corresponds to the optimal tree drawn from 1,000 bootstrap replicates. Isolates that belong to the B71 lineage are shown with orange (11 Zambian isolates) and red (77 Bangladeshi isolates and the Bolivian B71) background shades. **(B)** Geographical distribution of *M. oryzae* samples used in the tree. The colored points represent the approximate geographical origin of the isolates. Colors in (A) and (B) correspond to the plant host organism (upper inset). **(C)** Distribution of the collection year of *M. oryzae* wheat-infecting isolates used in the phylogenetic analysis.

We prioritized samples for whole genome sequencing based on our genotyping analyses and combined the samples with existing datasets to generate a set of 71 whole genome sequences of *M. oryzae* wheat isolates (*Triticum* lineage) from South America (N=39), Asia (N=21) and Africa (N=13) (Table S2). To gain insight into the phylogenetic relationship of the B71 lineage to other wheat isolates, we first performed unsupervised clustering of the 71 genomes using Principal component analysis (PCA) based on pairwise Hamming distances (Fig. 2A) and hierarchical clustering based on *f3*-outgroup statistics (Fig. S2). The B71 lineage shows reduced genetic diversity in comparison with South American isolates although incipient sub-structuring can be noted between Zambian and Bangladeshi clusters (Fig. 2A, inset).

**Fig. 2.**
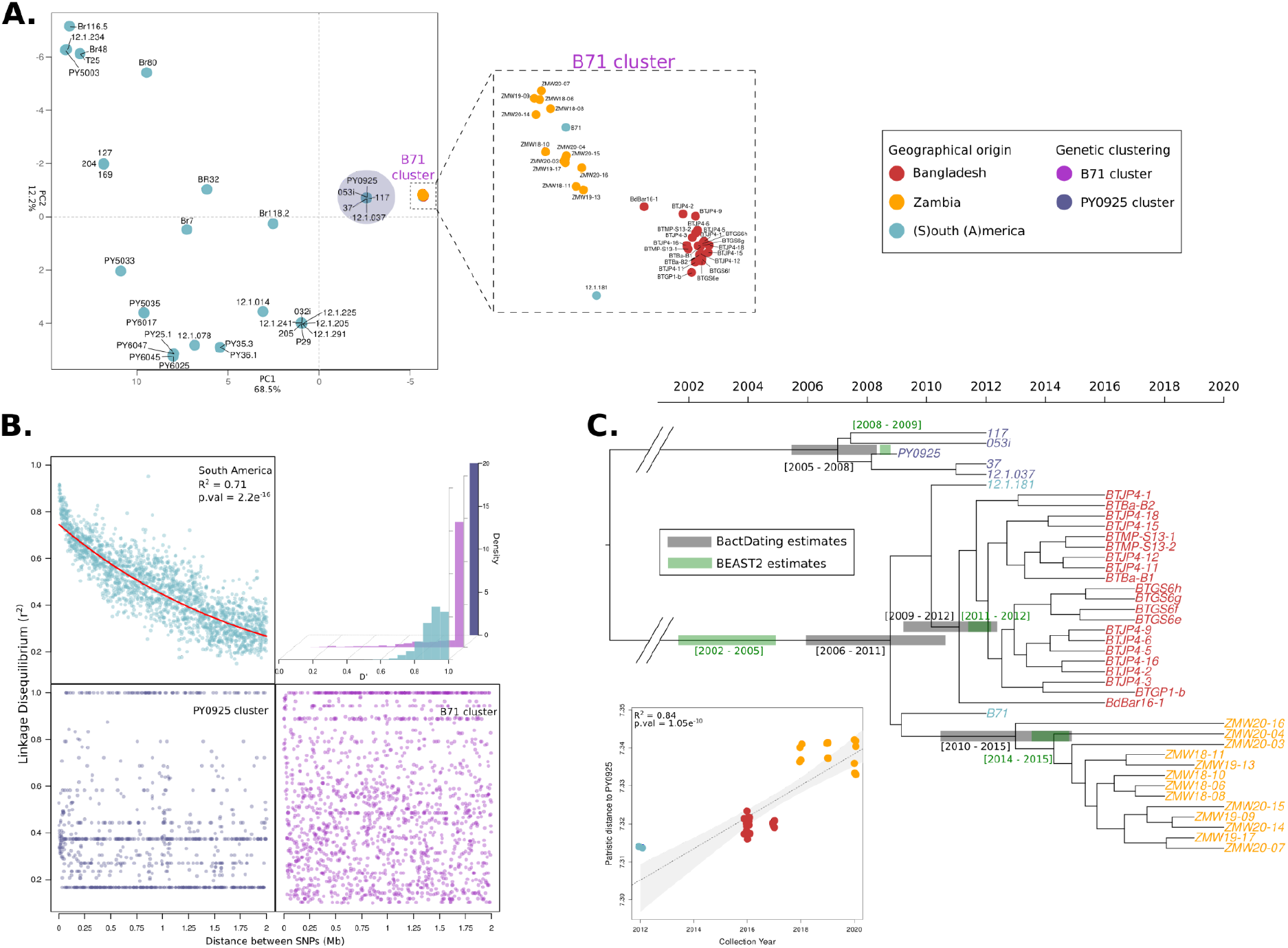
Wheat blast outbreaks in Zambia and Bangladesh originated by independent introductions of the recently emerged B71 clonal lineage. **(A)** The B71 lineage shows reduced genetic diversity in comparison with South-American wheat-infecting isolates. Principal component analysis (PCA) was performed based on genome-wide pairwise Hamming distances of 71 isolates from South America, Asia and Africa. The colors of the points indicate the provenance of each isolate (see inset). The circular shaded area indicates isolates from the Brazilian cluster (PY0925) that is the closest to the B71 cluster. Axes labels indicate the percentage of total variation explained by each PC. The magnified area shows isolates that are part of the B71 lineage. **(B)** The B71 cluster is a clonal lineage. The scatter plots show pairwise Linkage Disequilibrium (LD) (measured as *r*^*2*^) between SNPs that are at most two megabases apart. The red solid line in the South American cluster represents a fitted exponential decay model using nonlinear least squares. The histograms display LD expressed as *D’* for genome-wide SNPs. The points and bars are colored as indicated in the inset. **(C)** The B71 clonal lineage has recently expanded with independent introductions in Zambia and Bangladesh. The scatter plot shows the linear regression (dotted line) of root-to-tip patristic distances (y-axis) versus collection dates (x-axis) for the isolates of the B71 clonal lineage. Maximum likelihood tip-calibrated time tree of the B71 cluster isolates (the PY0925 cluster was used as an outgroup). The horizontal grey and green bars indicate 95% Confidence Intervals (CI) for divergence dates (in calendar years) calculated using BacDating and BEAST2, respectively. The points and isolate names are colored as indicated in the inset.

We decided to test the hypothesis that the B71 cluster is a clonal lineage and challenged it by measuring pairwise Linkage Disequilibrium (LD) (Fig. 2B; Fig. S3). Unlike the South American population, the B71 cluster displays no patterns of LD decay, which is consistent with clonality (Taylor et al. 2015; Adhikari et al. 2019) (Fig 2B, Fig. S3-S5).

We performed phylogenetic analyses to further define the genetic structure of the B71 clonal lineage. Owing to the much finer resolution obtained with genome-wide variation, we found that the Zambian and Bangladesh isolates clustered in separate well-supported clades with distinct phylogenetic affinities to South American isolates (Fig. 2C). We conclude that the clonal lineage has spread to Asia and Africa through two independent introductions, most probably from South America.

We leveraged the collection dates of *M. oryzae* clonal isolates to estimate their evolutionary rate. Before performing the tip-calibration analyses (Rieux and Balloux 2016), we removed regions that disrupt the clonal pattern of inheritance (Fig. S6-S8) (Didelot and Wilson 2015) and tested for a correlation between genetic distances and collection years (Fig. S9). We obtained rates ranging from 2.74e-7 to 7.59e-7 substitutions/site/year (Table S3). Using these rates, we dated the emergence of the Asian and African sub-lineage to similar periods (2009-2012 and 2010-2015, respectively) (Fig. 2C; Fig. S10). The B71 clonal lineage itself dates back to a few years earlier and probably emerged in South America around 2002-2011, before spreading to other continents (Fig. 2C; Fig. S10).

Can genome analyses inform practical disease management strategies of the wheat blast pandemic? Plant pathogens use secreted effector proteins to infect their hosts, but these effectors can also trigger plant immunity through an “avirulence” activity. The genome sequences of pandemic B71 lineage isolates offer the opportunity to identify effectors that can be targeted by the plant immune system. To date, the most predictive avirulence effector to confer wheat blast resistance is AVR-Rmg8 (Anh et al. 2015, 2018) which triggers an immune response in wheat carrying the resistance gene *Rmg8*. We scanned the available genomes and found that AVR-Rmg8 is conserved in all 71 isolates of the B71 clonal lineage even though the other 28 isolates of the *Triticum* lineage carry four diverse AVR-Rmg8 virulent alleles (eII, eII’, eII’’, eII’’’) that do not activate immunity (Fig. 3A; Fig. S11). B71 lineage isolates also lack the PWT4 effector, which is known to suppress AVR-Rmg8-elicited resistance (Horo et al. 2020; Inoue et al. 2021).

**Fig. 3.**
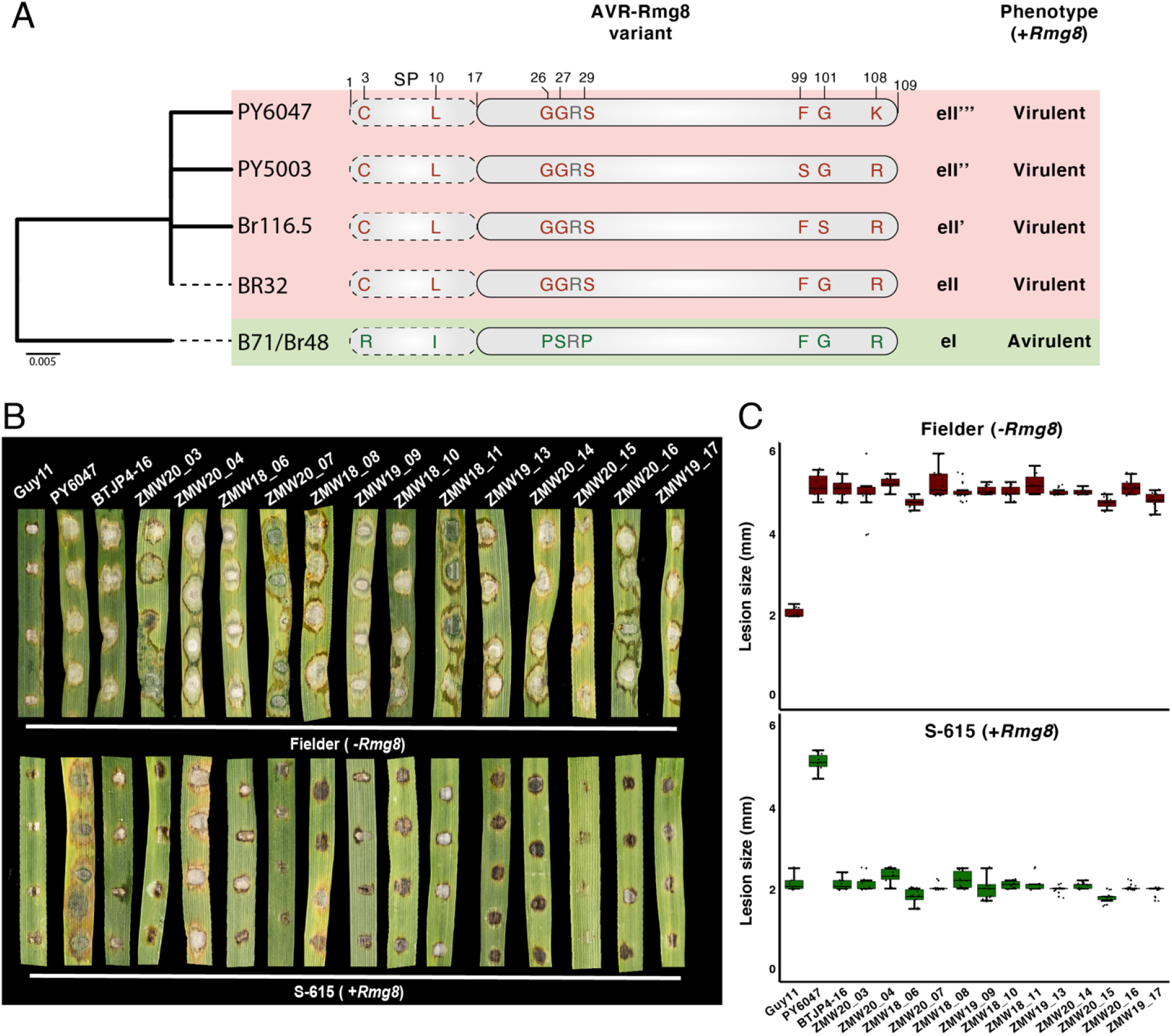
Rmg8 confers resistance against Zambian wheat blast isolates. **(A)** The wheat blast lineage contains five AVR-Rmg8 variants. Neighbour joining tree based on amino acid sequences of all non-redundant AVR-Rmg8 variants of 71 wheat blast lineage isolates (left). Representative isolate IDs are shown for each branch. Schematic representation of polymorphic amino acids in AVR-Rmg8 variants of the wheat blast lineage (center). Virulence phenotype associated with each AVR-Rmg8 variant on *Rmg8* containing host plants (right). **(B)** The resistance gene *Rmg8* is effective against wheat blast isolates collected in Zambia. Leaves from two weeks old seedlings of Fielder (-*Rmg8*, upper panel) and S-165 (+*Rmg8*, lower panel) wheat cultivars were inoculated with spores from Zambian wheat blast isolates, the rice blast isolate Guy11 (non-adapted, avirulent control), PY6047 (virulent control; AVR-Rmg8 eII’’’-carrier) and BTJP4-16 (avirulent on *Rmg8* carrying host plants, AVR-Rmg8 eI carrier). Disease and lesion size 5 days’ post-infection. **(C)** Quantification of lesions size (in mm) of 10 leaves and three independent experiments.

These genome analyses predict that the B71 lineage isolates (AVR-Rmg8 positive, PWT4 negative) cannot infect wheat plants with the matching disease resistance gene *Rmg8*. To test this, we inoculated 14 B71 lineage isolates from Zambia and Bangladesh on wheat lines with and without the *Rmg8* resistance gene (Fig. 3B; Fig. S12). Unlike a distinct South American isolate, none of these pandemic isolates could infect *Rmg8* wheat plants. We conclude that *Rmg8* is an effective resistance gene against the pandemic wheat blast lineage and has the potential to mitigate the spread of the disease.

Strobilurin fungicides are commonly used against the blast disease, but resistance is frequent in the South American population of the wheat blast fungus (Castroagudín et al. 2015). Genome analyses revealed that of the 71 wheat isolate genomes we examined, 13 carry the strobilurin resistance SNP (G1243C; Glycine to Alanine) in the mitochondrially-encoded Cytochrome B (*CYTB*) gene (Fig. 4A). Remarkably, all but one Brazilian isolate of the 36 B71 lineage genomes carry the G1243C allele and are predicted to be strobilurin sensitive. We tested this by assaying B71 lineage isolates and found that all tested 30 isolates are strobilurin sensitive (Fig. 4B-C; Fig. S13).

**Fig. 4.**
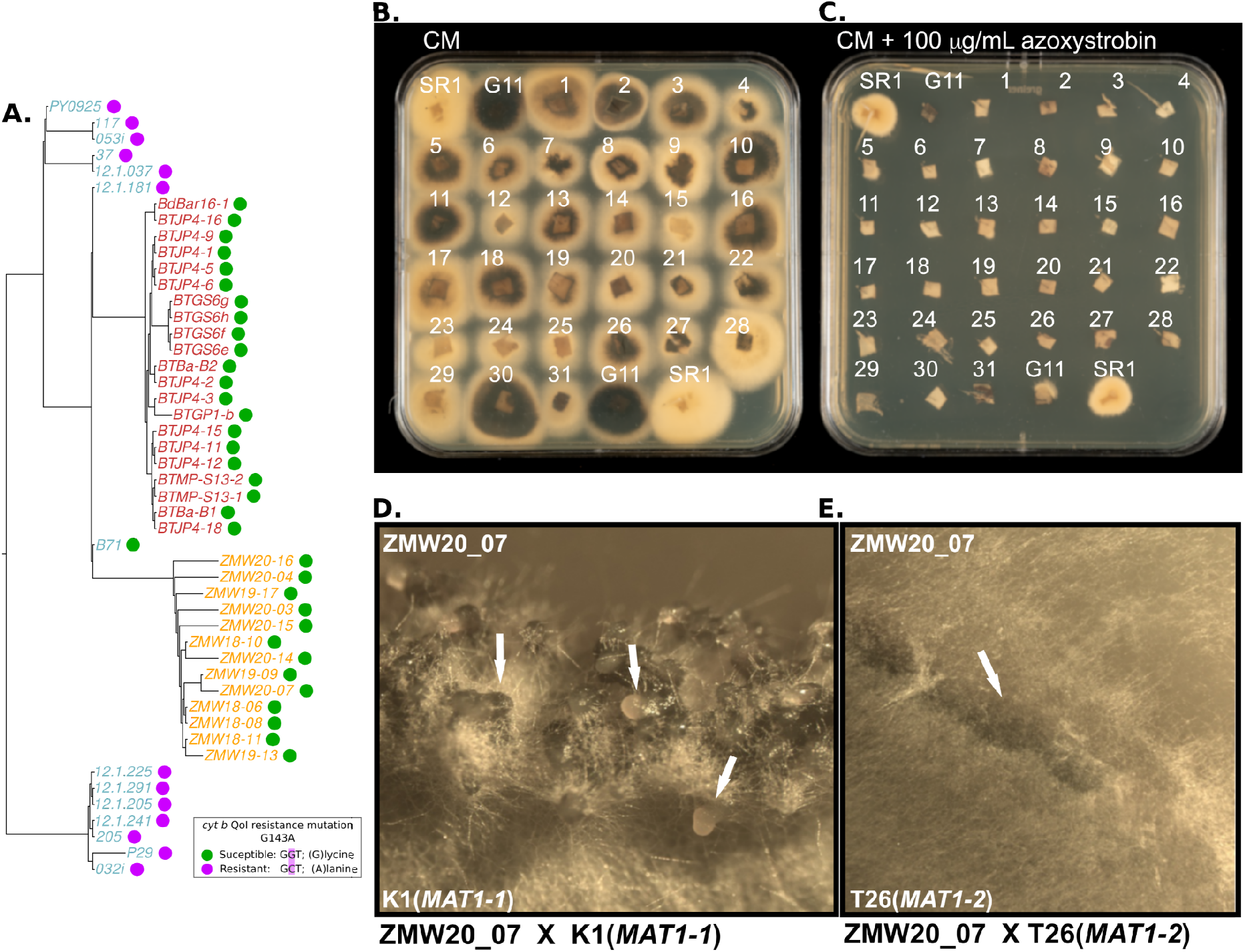
Zambian wheat blast isolates are susceptible to strobilurin fungicides but at risk from resistance development and can mate with prevailing finger millet blast isolates. (A) The tree describes, based on nuclear SNPs, the phylogenetic relationship among wheat-infecting blast isolates belonging to three clonal lineages: B71, PY0925 and P29. The tree was rooted in the midpoint. The coloured dots next to each isolate label represent the resistant-type allele of the mitochondrially-encoded *cyt b* gene associated with a susceptible or resistant predicted phenotype as shown in the inset. **(B-C)** 4×10^6^ spores of the Zambian strain ZMW20-14 were exposed to 100 μg ml^1^ azoxystrobin to obtain an azoxystrobin resistant strain with the G143A mutation (SR1). This strain, the rice infecting strain Guy11 (G11), and the wheat infecting strains numbered 1-31 respectively, BTBa-B1, BTBA-B2, BTGS6e, BTGS6f, BTGS6g, BTGS6h, BTGP1-b, BTMP-S13-1, PY6047, BTJP4-1, BTJP4-2, BTJP4-3, BTJP4-5, BTJP4-6, BTJP4-9, BTJP4-11, BTJP4-12, BTJP4-15, BTJP4-16, BTJP4-18, ZMW18-06, ZMW20-07, ZMW18-08, ZMW19-09, ZMW18-10, ZMW18-11, ZMW19-13, ZMW20-14, ZMW20-15, ZMW20-16 and ZMW19-17 were grown for 4 days at 25°C on CM in the absence (A) and presence (B) of azoxystrobin indicating that all the Zambian isolates have the ‘strobilurin susceptible’ genotype as anticipated by their *CYTB* sequences. **(D)** Zambian isolate ZMW20-7 (MAT-1-2) successfully produced perithecia when crossed with a finger millet blast isolate K1 (MAT-1-1) but **(E)** ZMW20-7 was unable to produce perithecia when crossed with a finger millet blast isolate T26 of the same mating type (MAT-1-2).

What is the evolutionary potential of the pandemic lineage of the wheat blast fungus? In laboratory experiments, we could readily recover spontaneous strobilurin (azoxystrobin) resistant mutants of African isolate ZMW20-14 (Fig. 4B-C) consistent with a high potential for emergence of fungicide resistance in the pandemic clonal lineage.

Multiple genetic lineages of the blast fungus are endemic to Africa causing destructive diseases on finger millet, rice and other grasses (Manyasa et al. 2019; Onaga et al. 2020). The spread of the B71 clonal lineage to Africa raises the spectre of sexual reproduction with endemic blast populations, which would further drive the evolutionary potential of the pandemic fungus. We determined that the pandemic B71 lineage belongs to the MAT1-2 mating type based on the genome sequences (Fig. S14) (Latorre, Were, et al. 2022), and therefore is predicted to mate with MAT1-1 isolates of *M. oryzae*. We tested and confirmed this prediction by showing that Zambian isolates from the pandemic lineage are fertile with MAT1-1 African finger millet isolates (Fig. 4D-E; Table S4).

The decade-old B71 clonal lineage of the wheat blast fungus, which spread twice from genetically diverse South American populations, happens to be avirulent on *Rmg8* wheat and sensitive to strobilurin fungicides. However, the emergence within short timescales of variants that are more damaging than the current genotypes is probable. This could happen either through mutations or sexual recombination with endemic blast fungus populations. Such variants could have increased virulence and fungicide tolerance thus adding to the difficulty in managing the wheat blast disease. These findings underscore the need for genomic surveillance to improve tracking and monitoring of the wheat blast fungus on a global scale and identifying variants of concern as soon as they emerge (World Health Organization 2022-2032).

Deployment of *Rmg8* wheat varieties on their own in affected areas is unlikely to provide sustained wheat blast management given that the fungus will probably evolve virulent races through AVR-Rmg8 loss of function mutations or gain of PWT4, an effector that suppresses the resistance conferred by Rmg8 (Inoue et al. 2021). *Rmg8* should therefore be combined with other resistance genes to be fully useful in areas with high fungal load and disease pressure. However, it might be judicious in the short-term to breed and deploy Rmg8 varieties in high-risk areas such as regions neighboring affected countries. This could serve as a firebreak to alleviate the need for wheat cultivation holidays as implemented in the Indian province of West Bengal, particularly in the face of likely wheat shortages and disruptions to global wheat trade (Mottaleb et al. 2019; Bentley 2022).

## MATERIALS AND METHODS

### Phylogenetic placement of *Magnaporthe oryzae* wheat-infecting isolates from the Zambian wheat blast outbreak (2018-2020) using a set 84 SNPs

To establish the genetic makeup and the phylogenetic placement of the wheat-infecting lineage that caused a wheat blast outbreak in Zambia (2018-2020) we analyzed a set of 84 SNPs, which were designed to distinguish between the pandemic clonal lineage of the wheat blast fungus (*Magnaporthe oryzae*) that reached South East Asia in 2016 from other *M. oryzae* genotypes (Islam et al. 2016). The dataset included 237 *M. oryzae* isolates from 13 different grasses that were genotyped using multiplex amplicon sequencing (“MonsterPlex”) (Tembo et al. 2021; Ascari et al. 2021). MonsterPlex is a highly accurate and cost-effective genotyping method to rapidly genotype field isolates (Tembo et al. 2021; Win et al. 2021). The Monsterplex genotypes were extracted from the genotyping matrices (Tembo et al. 2021; Ascari et al. 2021) as previously described (Latorre and Burbano 2021). To complement the MonsterPlex dataset and increase the geographic breadth of *M. oryzae* isolates, we extracted the 84 SNPs from 351 publicly available *M. oryzae* genomes. The joint dataset consisted of 537 worldwide distributed *M. oryzae* isolates (Fig. 1; Fig. S1; Table S1).

After removing adapters with *AdapterRemoval* v2 (Schubert, Lindgreen, and Orlando 2016), we mapped the Illumina-derived short reads to the *M. oryzae* B71 reference genome (Peng et al. 2019) using *BWA mem* (Li and Durbin 2009). To identify the genomic location of the 84 diagnostic SNPs in the *M. oryzae* B71 reference genome, we aligned each diagnostic SNP surrounded by 100 bp surrounding regions to the B71 reference genome using *blastn* (Altschul et al. 1990). Afterwards, for each of the per isolate mapped BAM files we retrieved the genotypes at each of the 84 genomic locations using *samtools mpileup* (Danecek et al. 2021). We concatenated all SNPs in a multi-fasta-like file that was used as input for phylogenetic analyses.

We built a Neighbour-Joining tree that includes a total of 532 *M. oryzae* isolates using the *R* package *phangorn* (Schliep 2011). We displayed a tree topology that corresponds to the optimal tree drawn from 1,000 bootstrap replicates (Fig. 1 and Fig. S1).

We further estimated the accuracy of the genotyping method by comparing SNP data acquired from 51 isolates using MonsterPlex to the SNPs extracted from matching genome sequences. We found that SNPs from both acquisition methods were identical whenever a SNP call was made (100%, 4,251/4,251 sites). In total, only 33 sites had gaps with missing data from MonsterPlex (0.77%, 33/4,284 sites) (Table S1).

### Processing of short reads and variant calling

Our phylogenetic analyses based on 84 SNPs (Fig. 1 and Fig. S1) confirmed our previous analyses, which showed that the emergence of wheat blast in Zambia and Bangladesh was caused by the same pandemic lineage of *M. oryzae* that traces its origin in South America (Latorre and Burbano 2021; Win et al. 2021). Consequently, from here on, we analyzed a set of 71 wheat-infecting *M. oryzae* genomes that belong to the wheat-genetic lineage. This set included isolates from South America (N=37), the outbreak in Bangladesh (2016-2018) (N=21), and isolates from the outbreak Zambia (2018-2020) (N=13) that were exclusively sequenced for this study (Tembo et al. 2021) (Table S2).

We removed adapters from the short reads from the set of 71 *M. oryzae* genomes using *AdapterRemoval* v2 (Schubert, Lindgreen, and Orlando 2016). To avoid reference bias, a common problem in population genetic analysis when comparing genetically similar populations, we mapped the short reads to the rice-infecting 70-15 reference genome (Dean et al. 2005) using *BWA mem2* (Li 2013; Vasimuddin et al. 2019) (Table S2). We used *samtools* (Danecek et al. 2021) to discard non-mapped reads, and *sambamba* (Tarasov et al. 2015) to sort and mark PCR optical duplicates.

As a first step to call variants, we used *HaplotypeCaller* from *GATK* (McKenna et al. 2010; DePristo et al. 2011) to generate genomic haplotype calls per individual using the duplicate-marked BAM files as input. Subsequently, we used *CombineGVCFs, GenotypeGVCFs* and *SelectVariants* from *GATK* (McKenna et al. 2010) to combine the individual genomic VCFs, call genotypes and filter SNPs, respectively. In order to select high quality SNPs to be included in our phylogenetic and population genetic analyses, we filtered SNPs using *Quality-by-Depth* (QD), which is one of the per-SNP summary statistics generated by GATK. Using *M. oryzae* genomes sequenced by two different sequencing technologies, we have previously shown that QD is the summary statistic that generates the best true positive and true negative rates (Latorre, Langner, et al. 2022). Following this approach, using *GATK VariantFiltration (McKenna et al. 2010)*, we kept SNPs with QD values no further away than one standard deviation unit away from the median of the distribution of QD values (Latorre, Langner, et al. 2022). Finally, we created a new VCF file keeping only non-missing positions using *bcftools* (Danecek et al. 2021).

### Population structure analyses

To assess the population structure of the 71 *M. oryzae* isolates we used genetic distances coupled with dimensionality reduction methods. First, we calculated pairwise Hamming distances using *Plink V*.*1*.*9* (Purcell et al. 2007). These distances were used as input for principal component analysis (PCA) using the function *prcomp* from the *R* package *stats* (R Core Team 2018) (Fig. 2A). Additionally, we used f3-outgroup statistics (Raghavan et al. 2014) to establish the pairwise relatedness between *M. oryzae* samples (X and Y) after divergence from an outgroup: *f3*(X, Y; outgroup). We used the rice-infecting *M. oryzae* 70-15 as an outgroup (Dean et al. 2005). We calculated *z*-scores for every possible pairwise sample comparison included in the f3-statistics test (*N* = 2,485). Subsequently, we carried out hierarchical clustering using the function *hclust* from the *R* package *stats* (R Core Team 2018) As input, we used a distance matrix generated from the f3-statistics-derived *f3* values (Fig. S2).

### Distinguishing clonality from outcrossing

To distinguish clonality from outcrossing in the B71 pandemic lineage and other genetic groups identified in our population structure analyses, we used patterns of Linkage Disequilibrium (LD) decay. While sexual reproduction (outcrossing) will generate patterns of LD decay that are driven by meiotic recombination, LD is not expected to decay in asexual non-recombining populations, i.e. the whole genome will be in LD. We analyzed LD decay patterns in the B71 lineage, the PY0925 lineage and treated the rest of Brazilians *M. oryzae* isolates as one group. To compute standard measures of LD we used *VCFtools* (Danecek et al. 2011). Since *VCFtools* is designed to handle diploid organisms, we transformed the haploid *VCF* files into “phased double haploid” *VCF* files. Afterwards, we computed pairwise LD as *r*^*2*^, and Lewontin’s *D* and *D’*. To evaluate LD decay, for each of the genetic groups, we calculated the average of each LD measure (*r*^*2*^, Lewontin’s *D* and *D’*) in bins of physical genomic distance (Fig. 2B and Fig. S3). To quantify the significance of LD decay, we fitted an exponential decay model using non-linear least squares. In order to compare the patterns of LD decay between the clonal lineages and the Brazilian group, we downsample the number of SNPs in the Brazilian group to the number of SNPs segregating in the B71 clonal lineage.

To perform a qualitative comparison between the observed patterns of LD decay with the expectations of LD decay in idealized populations we performed forward simulations using *FFPopSim* (Zanini and Neher 2012). We simulated genomes that consisted of 300 equidistant SNPs. We first sought to ascertain the effect of the probability of sexual reproduction per generation on the patterns of linkage disequilibrium decay. Each simulation was carried out for 100 generations keeping the population size, the crossover probability and the mutation rate constant, but changing the probability of sexual reproduction per generation (0,10^−3^,10^−2^,10^−1^ and 1) (Fig. S4). Additionally, we investigated the effect of the population size on the patterns of LD decay. To this purpose we simulated genomes that consisted of 200 equidistant SNPs. Each simulation was carried out for 100 generations keeping the crossover probability, the mutation rate and the probability of sexual reproduction per generation constant, but changing the population size parameter (10^2^,10^3^,10^4^,10^5^) (Fig. S5).

Finally, to detect recombination events in each of the genetic groups, we performed the four-gamete test (Hudson and Kaplan 1985) using *RminCutter* (Ross-Ibarra 2009). To be able to compare the number of violations of the four-gamete test among genetic groups, we normalized the number of violations of the four-gamete test by the number of segregating SNPs per genetic group (Fig. S6).

### Phylogenetic analyses, estimation of evolutionary rates and divergence times

To carry out phylogenetic analyses we used only the non-recombining genetic groups (clonal lineages) B71 and PY0925 (the latter was used as an outgroup) and included exclusively positions with no-missing data (full information). First, we generated a Maximum Likelihood (ML) phylogeny using *RAxML-NG* with a GTR+G substitution model and 1,000 bootstrap replicates (Kozlov et al. 2019). Then, we used the best ML tree as input for *ClonalFrameML* (Didelot and Wilson 2015), a software that detects putative recombination events and takes those into account in subsequent phylogenetic-based inferences. For every isolate we calculate the percentage of total SNPs masked by *ClonalFrameML* (Fig. S7). Additionally, we used pairwise Hamming distances to evaluate the levels of intra- and inter-outbreak genetic variation before and after *ClonalFrameML* filtering (Fig. S8).

Since the LD decay analyses revealed that the B71 pandemic lineage is a non-recombining clonal lineage, we hypothesized that the SNPs marked as putatively affected by recombination are preferentially located in genomic regions affected by structural variants, e.g. presence/absence variants. Such variants will generate phylogenetic discordances due to differential reference bias among the B71 isolates. To test this hypothesis we created full-genome alignments of the B71 and the 70-15 reference genomes using *Minimap2* (Li 2017) and visualized the output with *AliTV* (Ankenbrand et al. 2017). Then, we overlapped the visual output with the SNPs putatively affected by recombination that were previously identified by *ClonalFrameML* (Fig. S9).

To test for the existence of a phylogenetic temporal signal (i.e. a positive correlation between sampling dates and genetic divergence) we used the recombination-corrected tree generated by *ClonalFrameML* (Fig. 2C). Using this tree, and without constraining the terminal branch lengths to their sampling times, we measured tip-to-root patristic distances between all isolates of the B71 clonal lineage and the outgroup clonal lineage PY0925 with the *R* package *ape* (Paradis and Schliep 2019). We calculated the Pearson’s correlation coefficient between root-to-tip distances and sampling dates, and fitted a linear model using them as response and linear predictor, respectively (*distances ∼ sampling dates*) (Fig. 2C). We calculated 95% confidence intervals of the correlation signal between root-to-tip distance and collection dates by sampling with replacement and recalculating the correlation coefficient 1,000 times (Fig. 2C and Fig. S10). Additionally, to demonstrate that the obtained correlation coefficient was higher than expected by chance, we performed 1,000 permutation tests, where the collection dates were randomly assigned to the wheat blast isolates (Fig. S10).

To estimate the evolutionary rate and generate a dated phylogeny, where the divergence dates of all common ancestors are estimated, we used two different approaches. First, we used *BactDating*, a Bayesian framework that estimates evolutionary rates and divergence times using as an input a precomputed phylogenetic reconstruction (Didelot et al. 2018). As input for *BactDating*, we used the recombination-corrected tree generated by *ClonalFrameML*. The tree was loaded into *BactDating* using the function *loadCFML*, which permits the direct use of the output of *ClonalFrameML* as input for *BactDating* without the need of correcting for invariant sites (Fig. 2C). In our second approach we used *BEAST2*, a Bayesian approach, which in contrast to BactDating, co-estimates the evolutionary rate and the phylogenetic reconstruction (Bouckaert et al. 2014). From the alignment of the concatenated SNPs, we masked those that *ClonalFramML* marked as putatively recombining and used the masked alignment as input for the BEAST2 analyses. We carried out a Bayesian tip-dated phylogenetic analyses using the isolates’ collection dates as calibration points. Since, for low sequence divergence (<10%), different substitution models tend to generate very similar sequence distance estimates (Nascimento, Dos Reis, and Yang 2017), to simplify the calculation of parameters, we used the HKY substitution model instead of more complex models such as GTR. To reduce the effect of demographic history assumptions, and to calculate the dynamics of the population size through time, we selected a Coalescent Extended Bayesian Skyline approach (A. J. Drummond et al. 2005). The evolutionary rate was co-estimated using a strict clock model and a wide uniform distribution as prior (1E-10 - 1E-3 substitutions / site / year), which permits a broad exploration of the MCMC chains. To have genome-scaled results, the invariant sites were explicitly consider in the model by adding a “*constantSiteWeigths*” tag in the XML configuration file. Using the CIPRES Science Gateway (Miller, Pfeiffer, and Schwartz 2010), we ran four independent MCMC chains, each of which had a length of 20’000,000 with logs every 1,000 iterations. We combined the outputs with the *LogCombiner* and used *TreeAnotator* (Alexei J. Drummond and Bouckaert 2015; Bouckaert et al. 2014) to calculate the Maximum Credibility Tree as well as dating and support values for each node (Fig. 2C ; Fig. S11 ; Table S3).

### Determination of mating types

To assign the mating type for each isolate, we used two approaches. First, we created a fasta file containing the nucleotides codifying for the two mating type loci: MAT1-1-1 (GenBank: BAC65091.1) and MAT1-2-1 (GenBank: BAC65094.1) and used it as a reference genome. Using *bwa-mem2* (Li 2013; Vasimuddin et al. 2019), we mapped each isolate and used *samtools depth* (Danecek et al. 2021) to calculate the breadth of coverage for each locus as a proxy for the mating type assignment.

### Identification of AVR-Rmg8 effector variants and generation of the Avr-Rmg8 family tree

We used a mapping approach to identify Avr-Rmg8 family members in all 71 wheat blast lineage genomes. We mapped short reads sequencing data of each isolate to the B71 reference genome assembly (Peng et al. 2019) using BWA mem2 (Li and Durbin 2009) and extracted the consensus sequence of the AVR-Rmg8 locus from the output alignment files using *SAMtools* v.1.9 and *BCFtools* v.1.9 (Danecek et al. 2021) for each isolate. This led to the identification of five AVR-Rmg8 variants in 71 sequences.

### Generation of the Avr-Rmg8 family tree

To infer the AVR-Rmg8 family tree, we generated a multiple sequence alignment of 108 amino acid sequences of all identified variants using *MUSCLE* (Edgar 2004). We then removed pseudogenes and duplicated sequences to generate a non-redundant dataset and generated a maximum-likelihood tree with 1,000 bootstrap replications in *MEGA7* (Kumar, Stecher, and Tamura 2016).

### Leaf-drop and spray infection assay

To evaluate the response of Rmg8 against wheat blast isolates from Zambia, we carried out leaf drop and spray inoculations. An *Oryza*-infecting isolate from French Guyana, Guy11 was used as a negative control as this isolate does not infect wheat, while a *Triticum*-infecting isolate PY6047 from Brazil was used as a positive control it lacks the virulent allele of *AVR-Rmg8* (Jensen et al. 2019). Another *Triticum*-infecting Bangladesh isolate BTJP4-16 that carries an avirulent allele of *AVR-Rmg8* was also included. Wheat cultivars, S-615 (+Rmg8) and fielder (-Rmg8) were grown for 14 days in 9 cm diameter plastic plant pots or seed trays. Detached leaves from two weeks old seedlings were inoculated with fungal conidial suspension diluted to a final concentration of 1×10^5^ conidia mL-1 using drop inoculation method. The inoculated detached leaves were incubated in a growth room at 24°C with a 12h light period. Alternatively, plants were inoculated by spray infection using an artist’s airbrush (Badger. USA) as previously described in (Talbot, Ebbole, and Hamer 1993). After spray inoculation, the plants were covered in polythene bags and incubated in a negative pressure glasshouse with a 12 h light and dark cycle. Disease severity was scored after 5-6 days by evaluating lesion color and count or color and size for spray infection or drop inoculation, respectively. Lesions were classified as small black/brown non-spreading spots for resistant and large gray-spreading for susceptible. Each infection experiment was carried out three times.

### Generation of crosses and fertility status analysis

Generation of genetic crosses was carried as previously described in (Valent, Farrall, and Chumley 1991). A subset of Zambian isolates ZMW20_04, ZMW18_06, ZMW20_07, ZMW18_10, ZMW20_15 and ZMW20_16 (*MAT-1-2*) were tested against two finger millet tester isolates from Tanzania, T15 (*MAT-1-1*) or T26 (*MAT-1-2*), one from Kenya K1(*MAT-1-1*) and one from Ethiopia E12 (*MAT-1-1*). Two isolates were cultured opposite each other on oatmeal agar plate and incubated at 24 0 C for 7-10 days. The cultures were then transferred to a 20°C incubator until flask-shaped perithecia appeared at the crossing point. A Leica DFC360 FX microscope (Leica, Wetzlar, Germany) was used to visualize and image the formation of perithecia.

### Isolation of azoxystrobin resistant *Magnaporthe* strains

Isolation of azoxystrobin resistant *Magnaporthe* strains was carried out by exposure of spores of the fungus to azoxystrobin at 100g ml^-1^. Four million conidia of the Zambian wheat infecting isolate ZMW20-14 were recovered from CM grown agar cultures and plated on the surface of six 12 cm^2^ square Minimal medium agar plates (MM; (Foster, Jenkinson, and Talbot 2003)) with azoxystrobin at 100g ml^-1^. Plates were incubated for 2 weeks and then overlaid with MM plus 100ug ml^-1^ azoxystrobin and grown for a further week. Five colonies were isolated from these plates and colonies were purified by subculture to MM with 100ug ml^-1^ azoxystrobin for a further week before genotyping. PCR competent genomic DNA was isolated from a 4 mm^2^ plug of mycelium from the purified azoxystrobin resistant colonies (named AZ1-AZ5) with disruption using a Geno/grinder ® SPEX SamplePrep. To extract the mycelial plug it was placed in 400 l of extraction buffer (1M KCl, 10 mM Tris-HCl (pH 8), 5mM EDTA (pH 8) together with a 5 mm stainless steel bead (Qiagen) and macerated at 1,200 rpm for 2 mins in the Geno/grinder followed by a 10-minute centrifugation (17,000 x g) in a bench top centrifuge at room temperature. 300l of the supernatant was transferred to a new tube and nucleic acids were precipitated with 300 l of isopropanol followed by a 10-minute centrifugation at room temperature at 17,000 x g. Nucleic acid pellets were air dried for 10 minutes and then resuspended in 50 ul of TE with 50 μg ml^-1^ RNAse by vortexing and then incubated for 10 minutes at 37°C. 1 l of the DNA was used in a 50 l PCR reaction with the enzyme Q5 polymerase (New England Biolabs) and the primers Cytb-f AGTCCTAGTGTAATGGAAGCand Cytb-r ATCTTCAACGTGTTTAGCACC (annealing temperature 61.5°C). Amplicons were sequenced following gel purification in both directions using the same primers used to generate them to score for the presence or absence of the strobilurin resistance conferring mutation (Castroagudín et al. 2015).

### Datasets and preliminary reports

Genotyping and whole-genome sequencing data included in this article were released without restrictions as soon as they were produced through the OpenWheatBlast Community (https://zenodo.org/communities/openwheatblast). OpenWheatBlast collects research output datasets on wheat blast and encourages scientists to analyze and share them before formal publication. We list below the preprints that were shared through the OpenWheatBlast community and whose data were analyzed in this publication:

## Availability of code and data

The datasets and scripts generated during and/or analyzed during the current study are available in the Github repository: https://github.com/Burbano-Lab/wheat-clonal-linage

## Acknowledgments

We thank Aida Andrés, members of her group at UCL and Michael Dannemann for input on data analyses, and Talia Karasov for comments on the manuscript. We also thank Emilie Chanclud, as well as Emerson M. Del Ponte and group for contributions to the genotyping experiments.

## Funding

This project was funded by:

- The Leverhulme Trust (Philip Leverhulme Prize) (HAB).
- Royal Society (RSWF\R1\191011) (HAB).
- Gatsby Charitable Foundation (The Sainsbury Laboratory). - BBSRC BBS/E/J/000PR9795, BBS/E/J/000PR9796, BBS/E/J/000PR9798, BB/P023339/1, BB/W008157/1 (The Sainsbury Laboratory).
- BBSRC equipment grant (BB/R01356X/1) (University College London).
- The Krishi Gobeshona Foundation (KGF) of Bangladesh grants Nos. KGF TF50-C/17 and TF 92-FNS/21 (TI).
- European Research Council BLASTOFF (SK).

## Author contributions

*Conceptualization and experimental design:* SML, VMW, AJF, TL, AM, AH, CJ, JW, NT, HAB, SK.

*Experiments:* SML, VMW, AJF, TL, AM, AH, CJ, LR, DMS, DC, KHL.

*Data analysis:* SML, VMW, AJF, TL, AM, AH, SR, CJ, DMS, SGOS, YT, DC, KHL, JW, NT, HAB, SK.

*Contribution of materials or analysis tools:* SML, VMW, AJF, TL, SA, SR, DRG, WM, NUM, MSM, RMM, ANMM, SKP, SMFR, AAMR, LR, RS, SS, PKS, AB, YT, DC, KHL, TI, BT, JW.

*Writing:* SML, VW, AJF, TL, JW, NT, HAB, SK.

## Competing interests

- KHL is a founder of Floodlight Genomics.
- TI receives funding from Krishi Gobeshona Foundation of Bangladesh.
- SK receives funding from industry and has filed patents on plant disease resistance.

## Notes

https://github.com/Burbano-Lab/wheat-clonal-linage.git

### Competing Interest Statement

KL is a founder of Floodlight Genomics.
TI receives funding from Krishi Gobeshona Foundation of Bangladesh
SK receives funding from industry and has filed patents on plant disease resistance.

