## Supplemental Figures for "A pandemic clonal lineage of the wheat blast fungus"

### SUPPLEMENTARY FIGURES

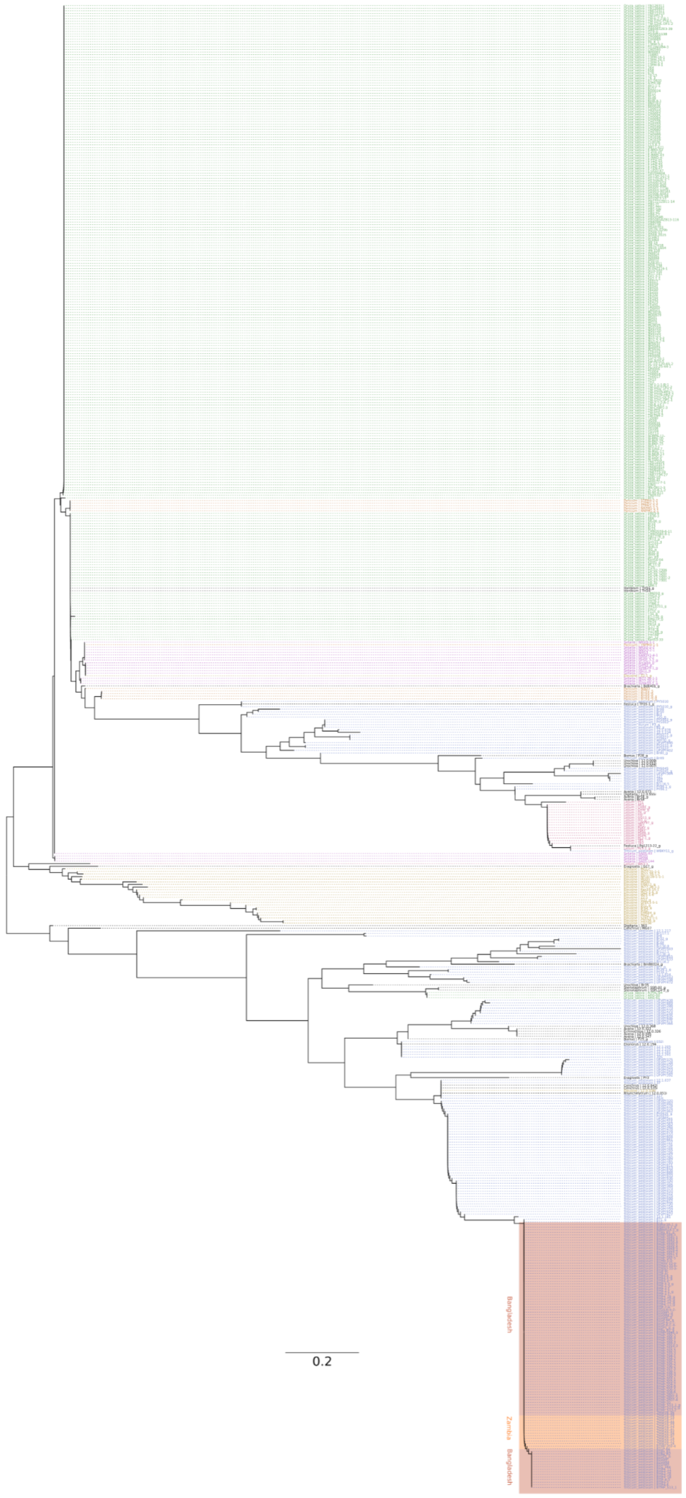


**Fig. S1. The emergence of wheat blast in Bangladesh and Zambia was caused by the B71 genetic lineage of *Magnaporthe oryzae*.** Neighbor-joining tree of 576 worldwide distributed blast isolates based on 84 concatenated SNPs. The topology corresponds to the optimal tree drawn from 1,000 bootstrap replicates. Names of host organisms are shown together at the tips. Duplicated identifiers without the "_*g*" suffix, represent data belonging from the same isolate but from the "monsterplex" set of 84 SNPs.

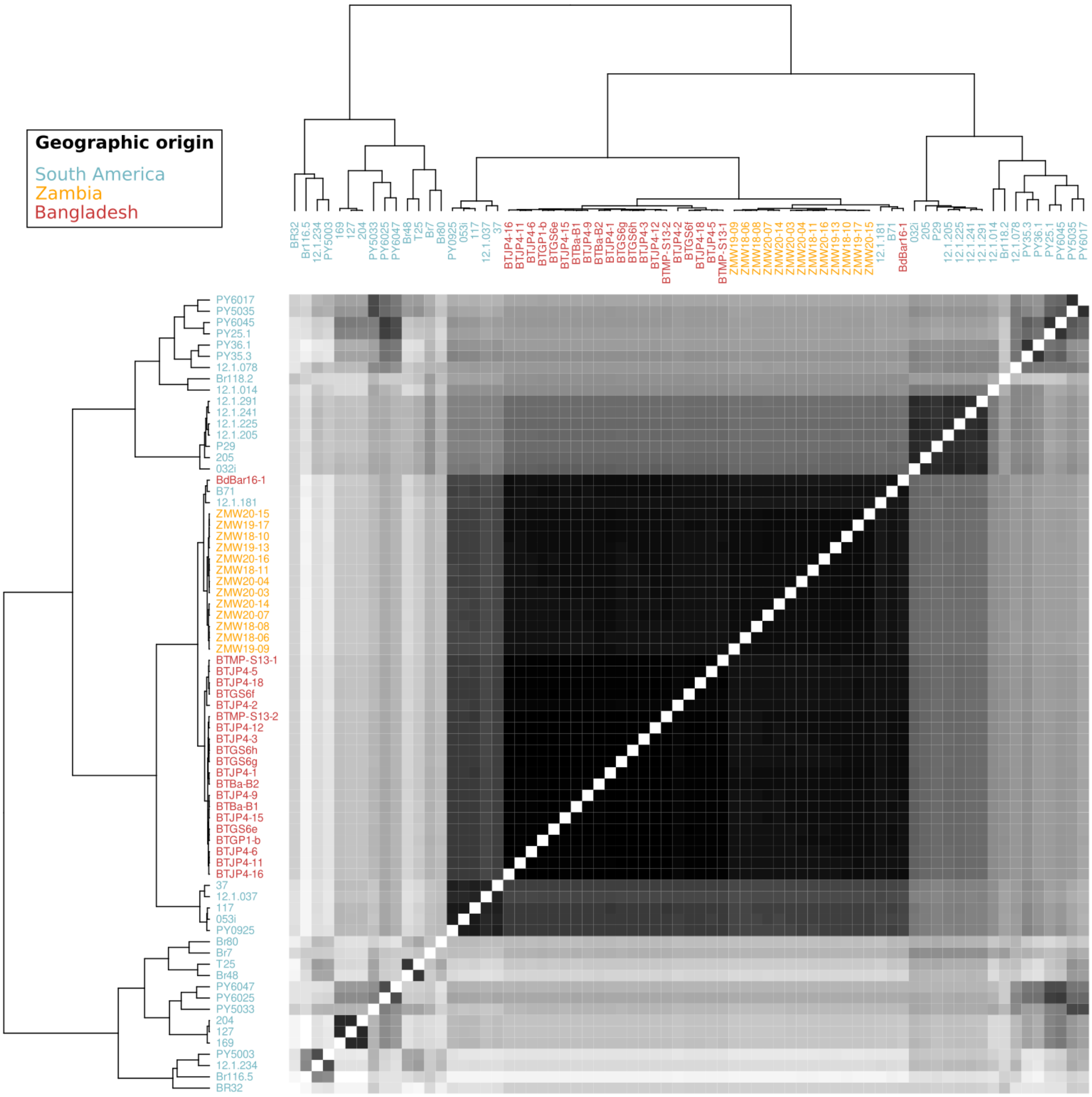


**Fig. S2. Genetic clustering of *Magnaporthe oryzae* identifies isolates from the Bangladesh and Zambian outbreaks as part of the B71 lineage.** The pairwise relatedness between *M. oryzae* samples (X and Y) was estimated using *f3*-outgroup statistics of the form *f3*(X, Y; outgroup), which measures the amount of shared genetic history (genetic drift) between X and Y after the divergence from an outgroup (rice-infecting *M. oryzae* isolate 70-15). The hierarchical clustering is based on *f3*-scores resulting from *f3*-outgroup statistic calculations. Darker colors indicate more shared drift.

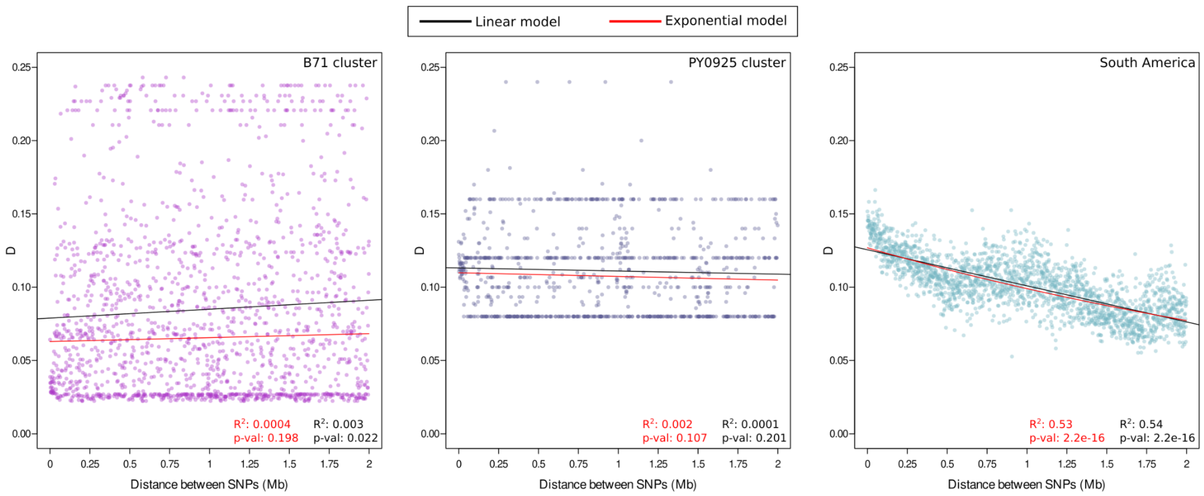


**Fig. S3. Linkage Disequilibrium (LD) does not decay with physical distance in wheat-infecting blast isolates from the B71 lineage**. The scatter plots show pairwise Linkage Disequilibrium (LD) (measured as *D*) between SNPs that are at most two megabases apart. The solid lines represent fitted linear and exponential decay models as indicated in the inset.

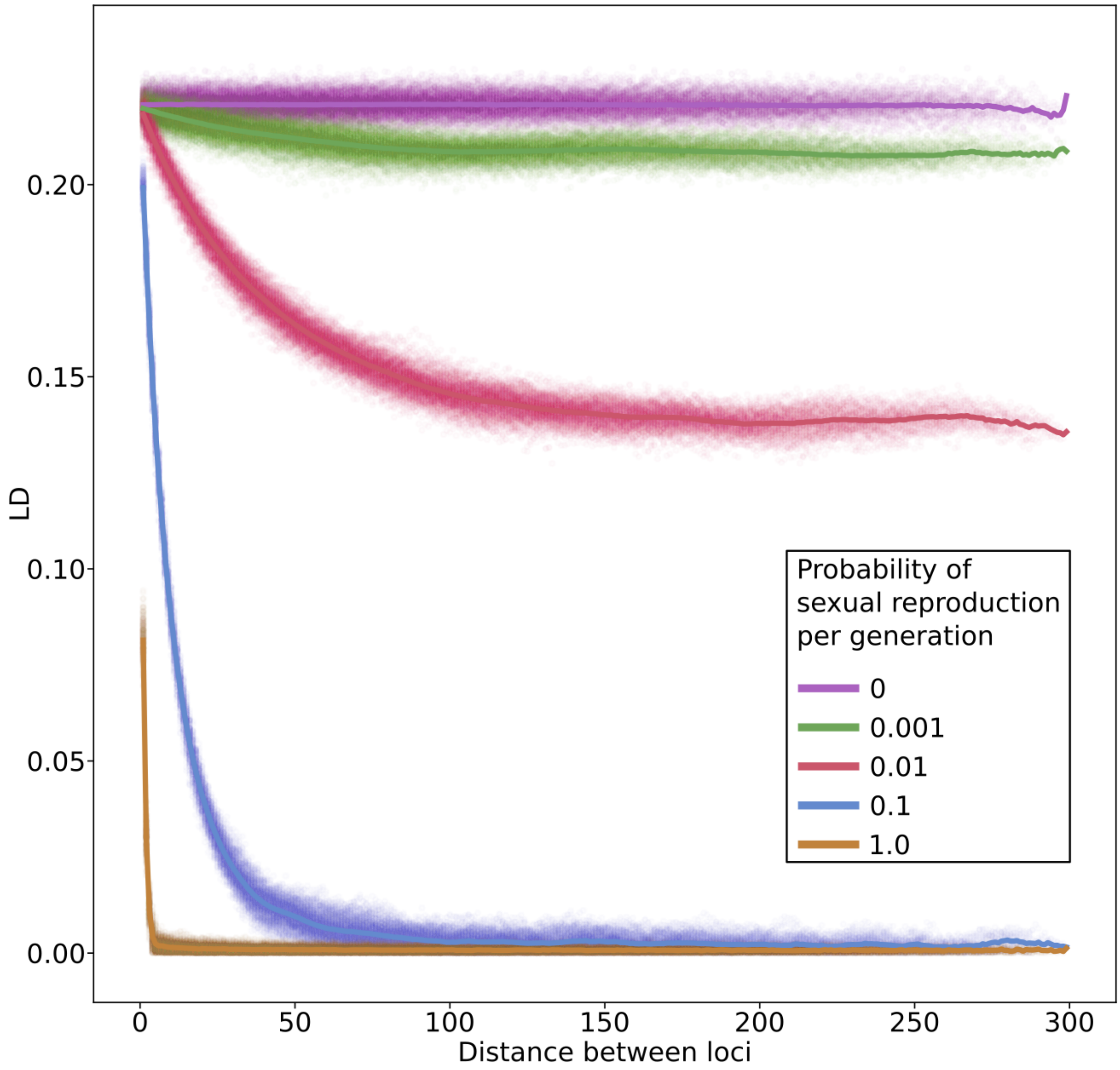


**Fig. S4. Forward simulations indicate that the probability of sexual reproduction per generation determines the extent of Linkage Disequilibrium (LD) decay**. The simulated genomes consisted of 300 equidistant SNPs. Each simulation was carried out for 100 generations keeping the population size, crossover probability and the mutation rate constant, but changing the probability of sexual reproduction per generation (see inset). Dots represent LD (measured as *D*) as a function of the distance between two loci and thick lines represent the mean value per distance-bin. Points and lines are coloured as indicated in the inset.

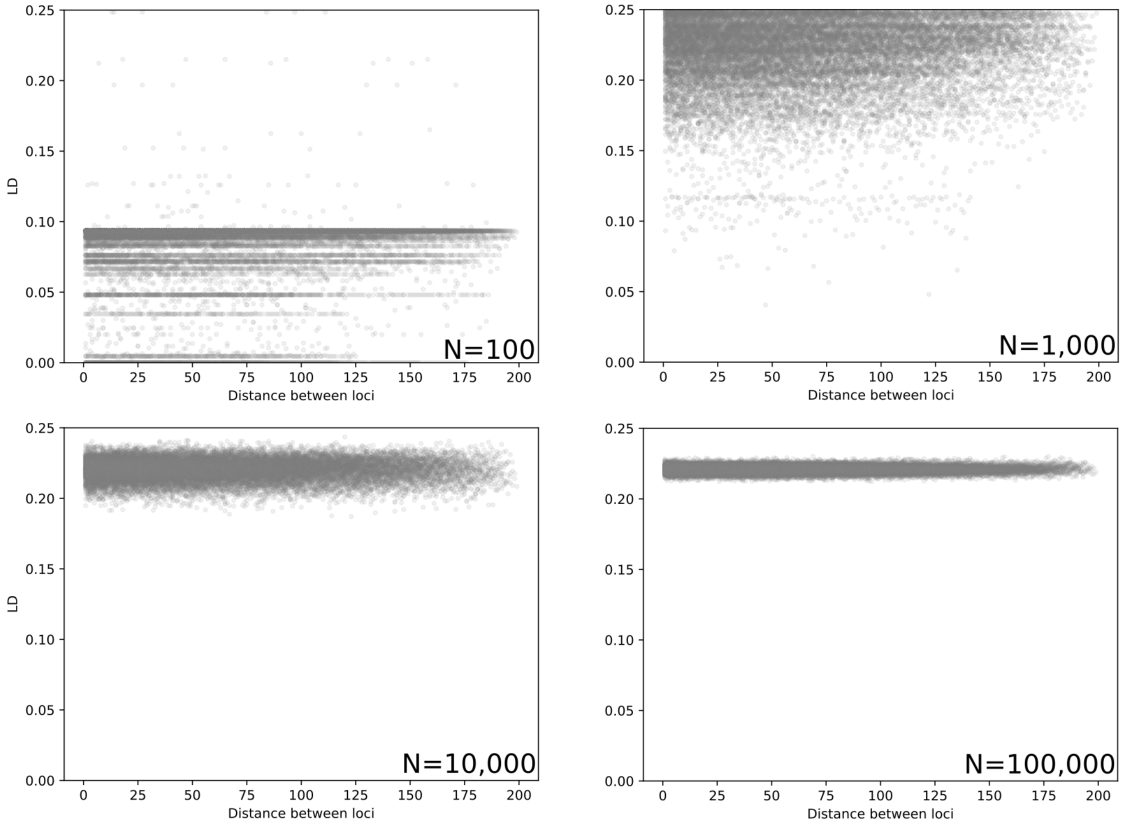


**Fig. S5. Forward simulations indicate that LD breaks as a function of population size**. The simulated genomes consisted of 200 equidistant SNPs. Each simulation was carried out for 100 generations keeping the crossover probability, the mutation rate and the probability of sexual reproduction per generation constant, but changing the population size parameter. Dots represent LD (measured as *D*) as a function of the distance between two loci.

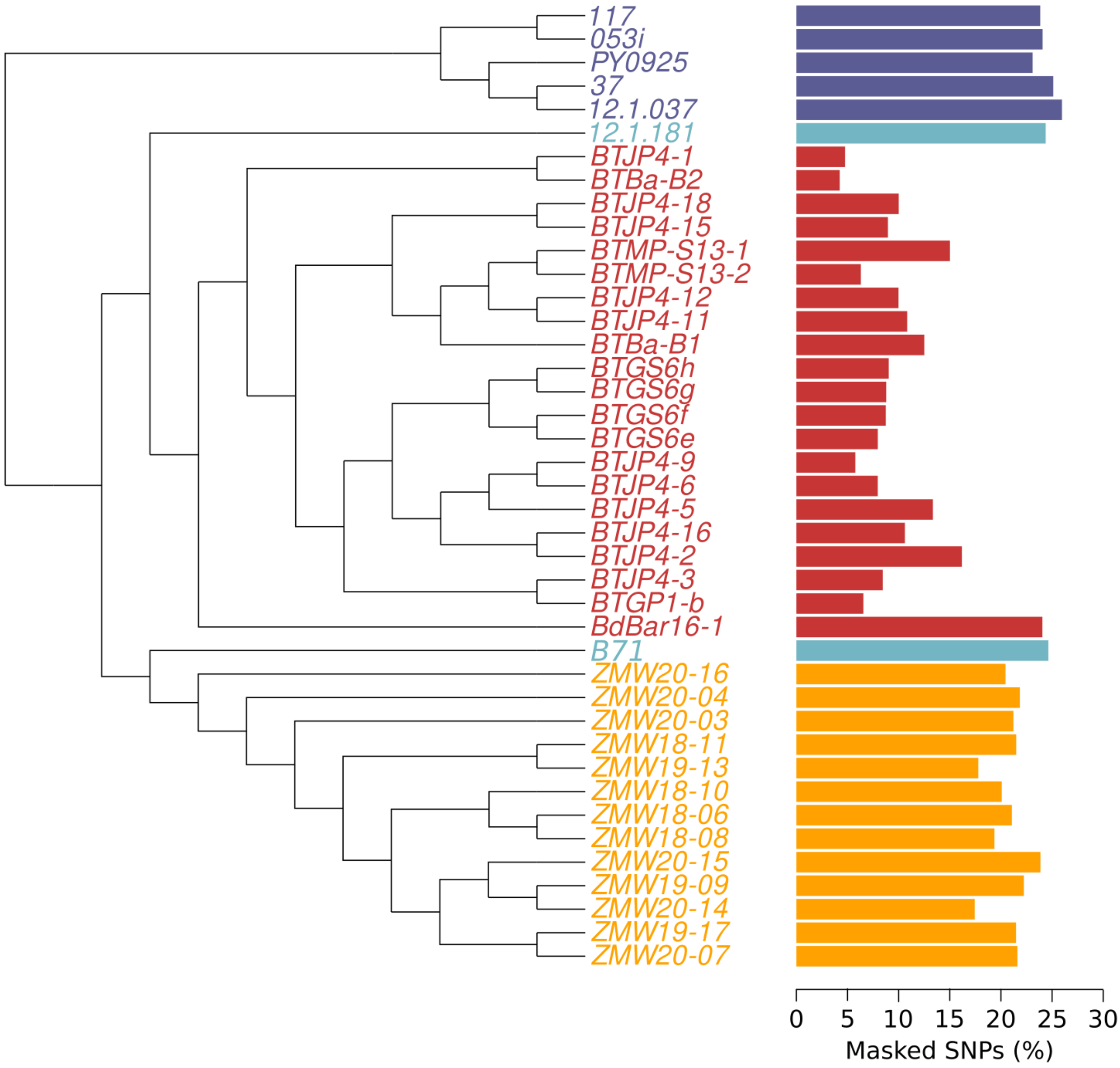


**Fig. S6. *ClonalFrameML* identifies on average 16% of SNPs/per isolate as putatively recombining**. The dendrogram shows the phylogenetic relationships of *Magnaporthe oryzae* strains as inferred by *RAxML-NG*. The dendrogram is schematic, i.e. the branch lengths do not accurately represent phylogenetic distances. The bars show the percentage of SNPs identified as putatively recombining by *ClonalFrameML*, which were masked in our dating analyses. The bars and isolate names are colored as indicated in the inset.

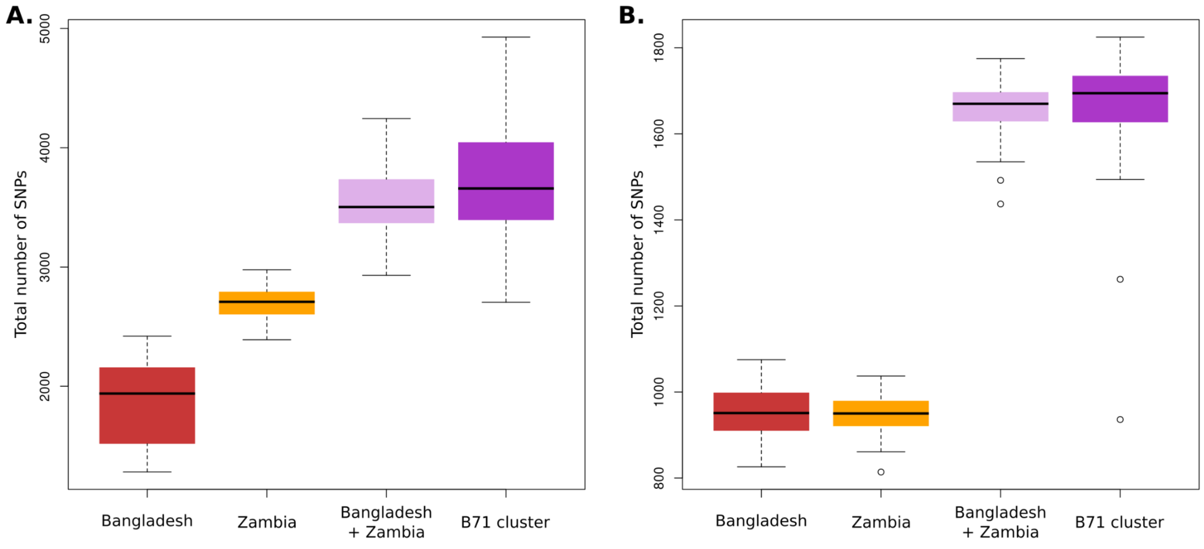


**Fig. S7. The outbreaks of Bangladesh and Zambia show similar levels of genetic diversity**. The panels show the total number of segregating SNPs in the outbreaks of Zambia, Bangladesh and the B71 clonal lineage. **(A)** Total number of segregating SNPs. **(B)** Total number of SNPs after excluding putatively recombining SNPs identified *ClonalFrameML*. All groups include 13 isolates that were sampled with replacement 100 times.

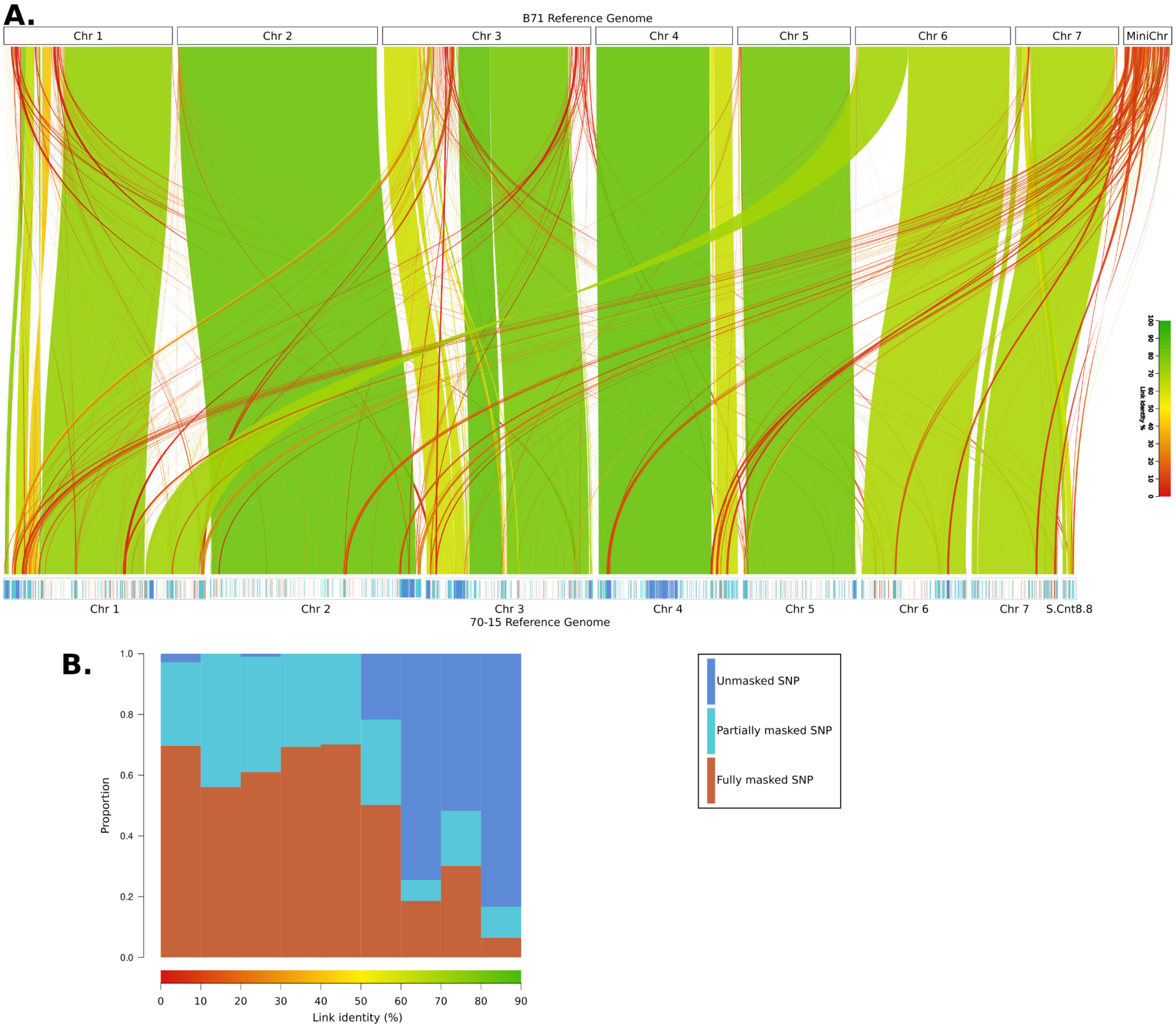


**Fig. S8. Putative recombinant regions are likely caused by structural variation**. **(A)** The upper horizontal track is a representation of the wheat blast B71 reference genome. The bottom horizontal track is a representation of the 70-15 rice blast reference genome. Vertical ticks represent different types of SNPs (dark blue: unmasked SNPs; light blue: partially masked SNPs, i.e. masked in only some samples; and red: SNPs masked in all samples) (inset). Unmasked and partially masked SNPs were included in the phylogenetic analyses, whereas fully masked SNP were excluded from them. Ribbons represent syntenic regions between the rice blast 70-15 genome and the wheat blast B71 reference genomes. The coloring of the ribbons indicates the level of identity (chromatic scale). **(B)** The barplot summarizes the proportion of the three SNP categories (inset) per bin of link identity between the B71 and 70-15 reference genomes.

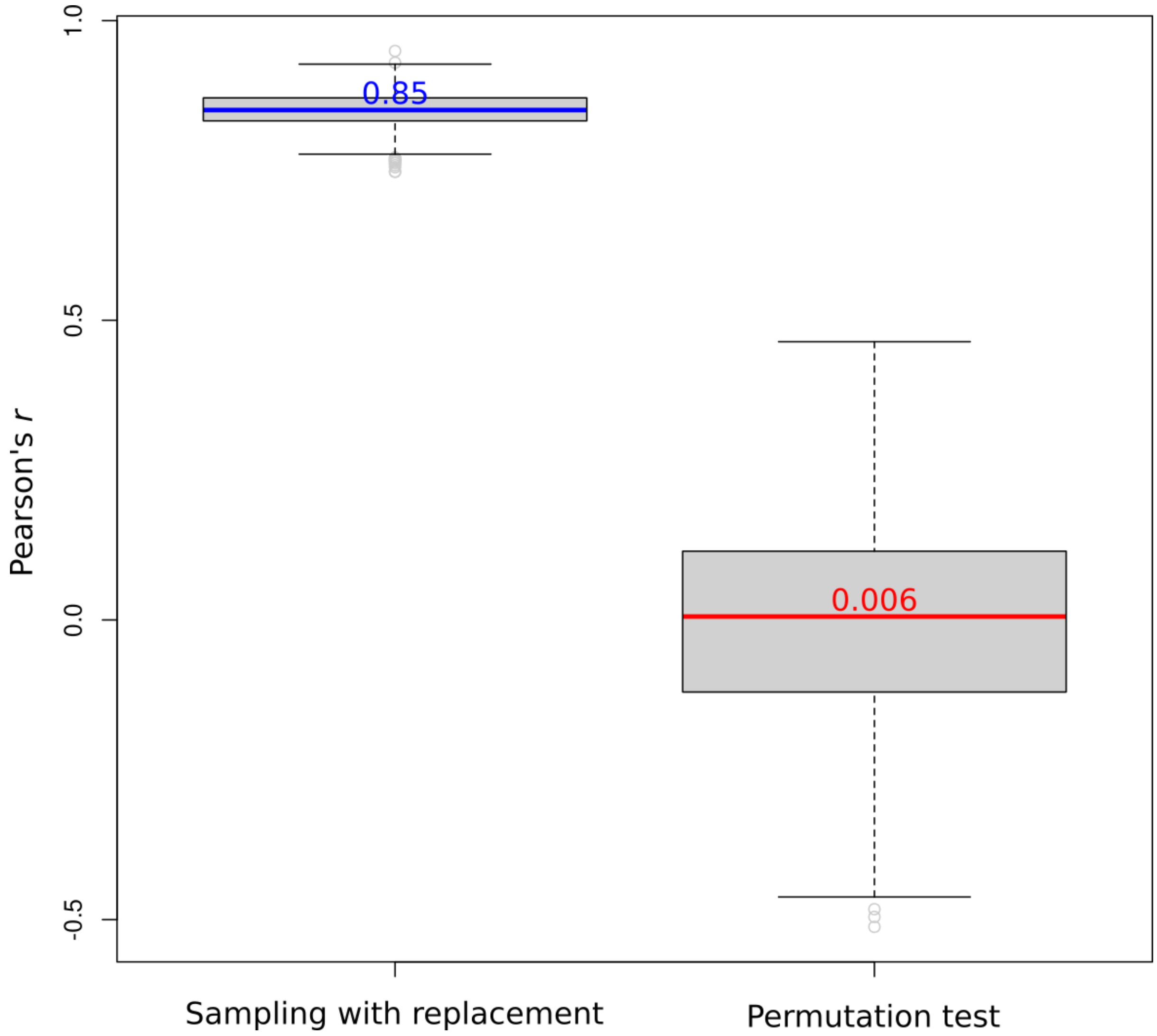


**Fig. S9. The temporal signal of the B71 lineage is robust and significantly bigger than expected by chance**. Boxplots represent the distribution of the Pearson's *r* correlation between root-to-tip patristic distances and collection dates of the isolates of the B71 clonal lineage. The left boxplot depicts the distribution of 1,000 instances of sampling with replacement from the original dataset. The right boxplot represents the distribution of 1,000 permutation tests, where collection dates were randomly assigned to wheat blast isolates. Median values are shown within each boxplot.

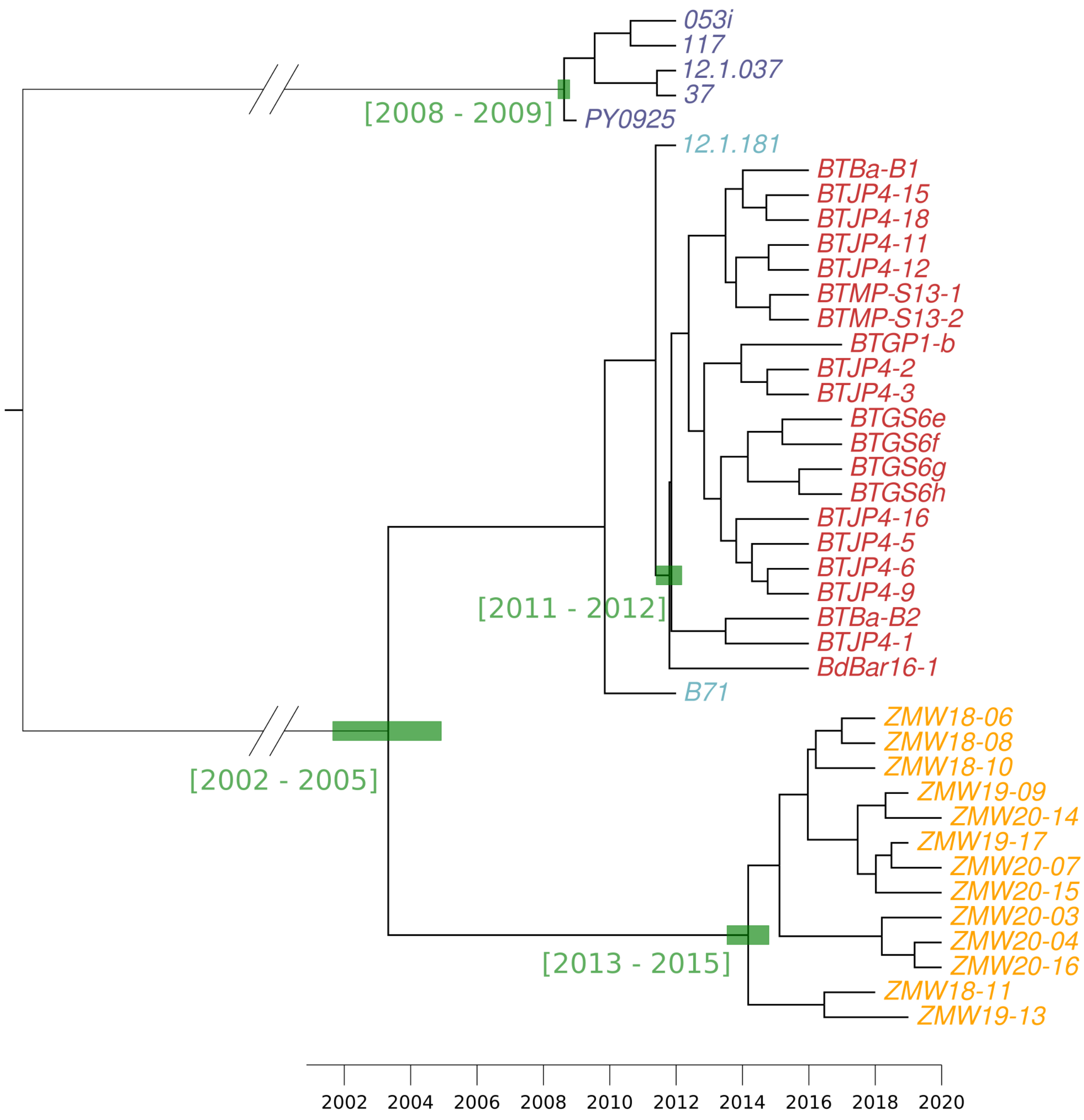


**Fig. S10. Bayesian tip calibrated phylogenetic tree using individuals belonging to B71 and PY0925 clonal lineages**. Average, and HPD 95% confidence intervals are shown in calendar years for nodes leading to the outbreaks or clonal lineage expansions.

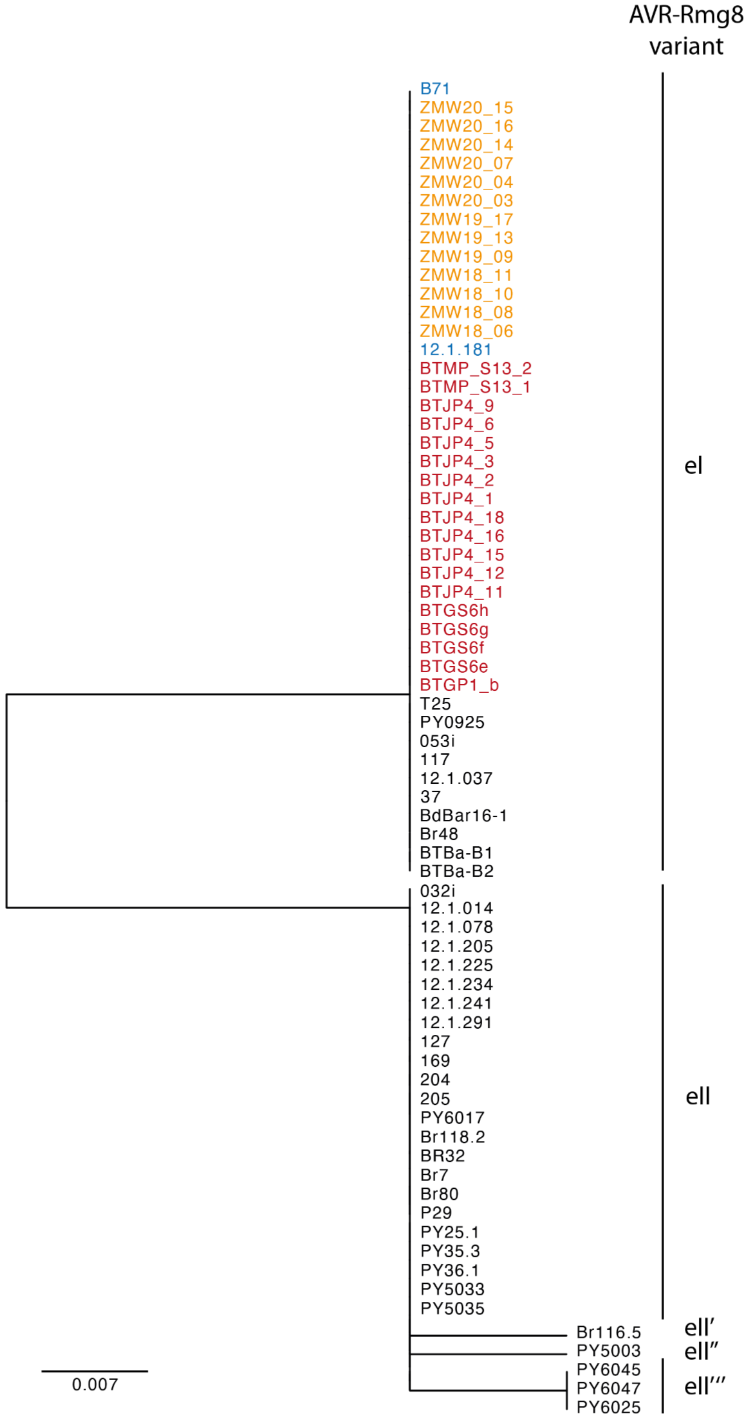


**Fig. S11. The pandemic wheat blast lineage contains the avirulent AVR-Rmg8 variant eI.** Neighbour joining tree based on amino acid sequences of all AVR-Rmg8 variants of the wheat blast lineage reveals diversifying AVR-Rmg8 alleles. Isolate IDs are shown for each branch. Wheat blast isolates of the pandemic B71 lineage are colored.

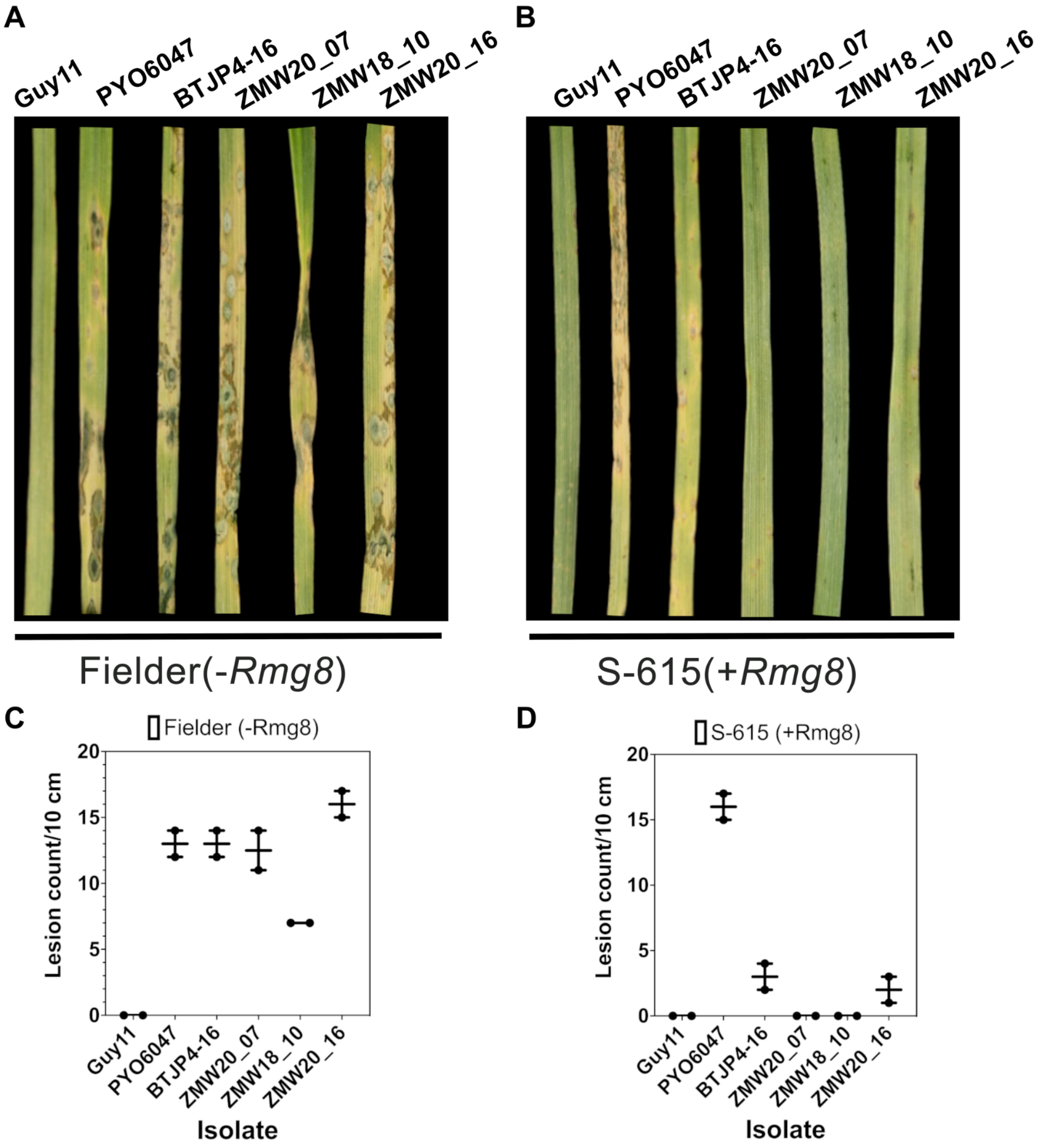


**Fig. S12. Rmg8 confers resistance against the Zambian wheat blast population**. **(A)** Leaves from two weeks old seedlings of Fielder (-Rmg8) and **(B)** S-165 (+Rmg8) wheat cultivars were inoculated with spores from ZMW20_07, ZMW18_10, ZMW20_16, Guy11, PY6047 and BTJP4-16 using a spray infection method. Disease lesions were scored 6 days' post-infection. **(C-D)** Boxplots show lesion count per 10 cm for two independent experiments.

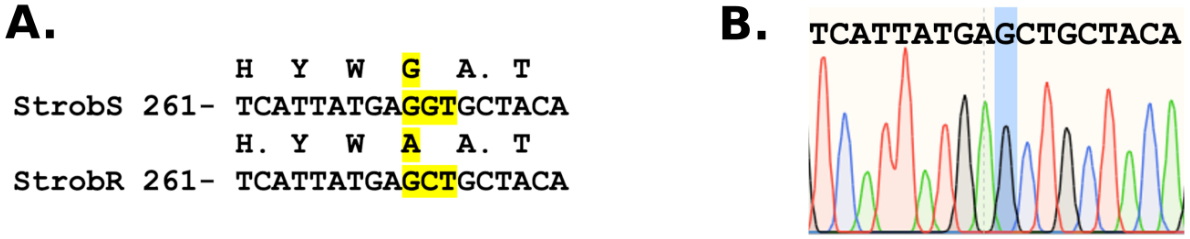


**Fig. S13. All Zambian and Bangladeshi wheat-infecting blast isolates are susceptible to strobilurin fungicides**. **(A)** The 70 wheat blast isolates had just two genotypes with respect to the mitochondrially encoded gene *CYTB*. The GGT to GCT mutation in the *CYTB* gene results in a substitution at position 143 in the gene product and is known to confer resistance to strobilurin class fungicides. All the Zambian isolates sequenced have the ‘strobilurin susceptible’ genotype. **(B)** Sequencing of the *CYTB* partial gene sequence in the azoxystrobin resistant strain (SR1) indicated a homogenous population of mitochondria with the CytB G143A genotype.

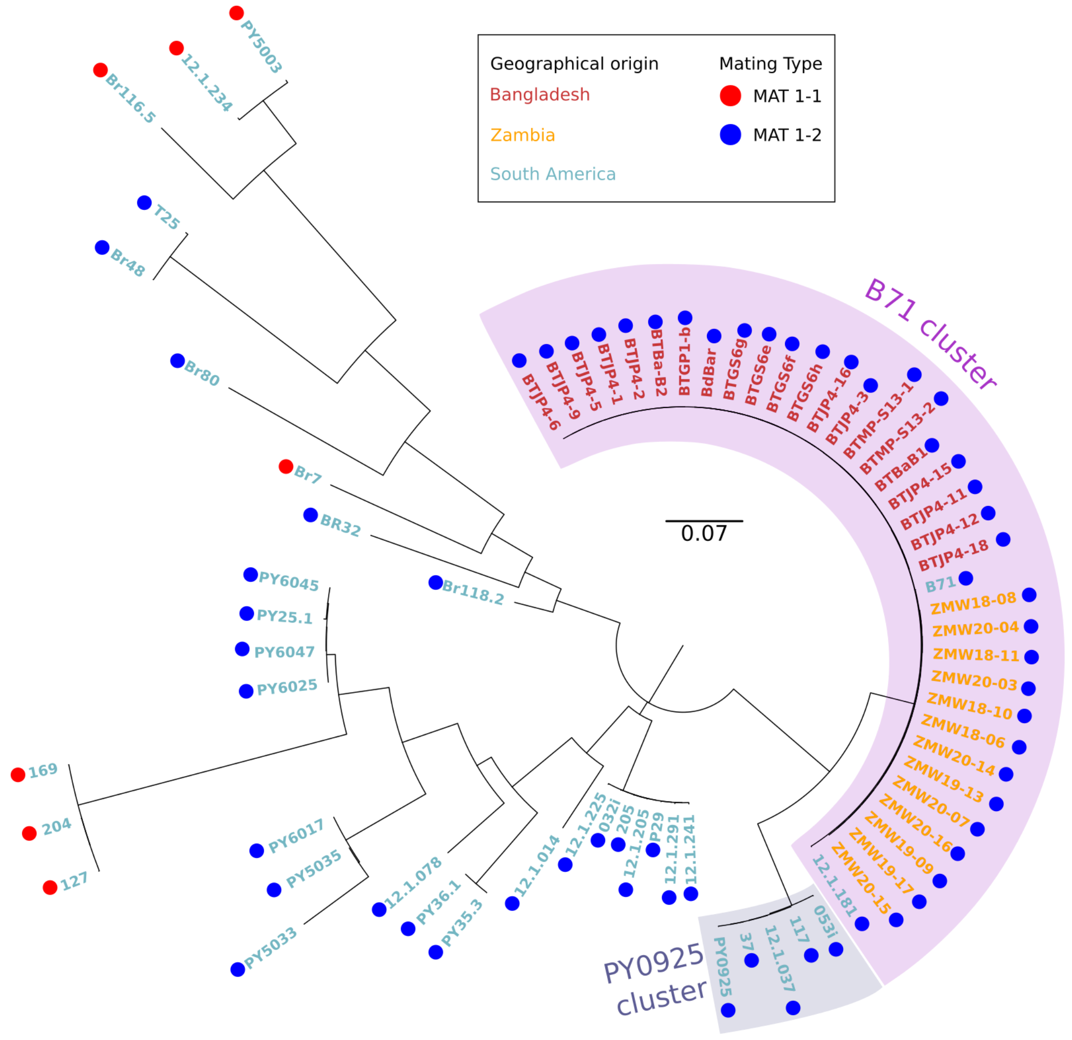


**Fig. S14. Only the MAT 1-2 segregates in the B71 clonal lineage**. The Neighbor-Joining tree describes the phylogenetic relations among wheat-blast infecting isolates. The mating types are codified as colored circles next to the isolate name (see inset).
